# TLR7 ligation augments haematopoiesis in Rps14 (uS11) deficiency via paradoxical suppression of inflammatory signalling and enhanced differentiation

**DOI:** 10.1101/2020.07.06.175000

**Authors:** Oscar A Peña, Alexandra Lubin, Jasmine Rowell, Catherine Hockings, Youngrock Jung, Yvette Hoade, Phoebe Dace, Leonardo E Valdivia, Karin Tuschl, Charlotta Boiers, Maria C Virgilio, Simon Richardson, Elspeth M Payne

## Abstract

Myelodysplastic syndrome (MDS) is a haematological malignancy characterised by blood cytopenias and predisposition to acute myeloid leukaemia (AML). Therapies for MDS are lacking, particularly those that impact the early stages of disease. We developed a model of MDS using zebrafish using knockout of Rps14, the primary mediator of the anaemia associated with del (5q) MDS. These mutant animals display dose- and age-dependent abnormalities in haematopoiesis, culminating in bone marrow failure with dysplastic features. We utilized rps14 knockdown to undertake an *in vivo* small molecule screen to identify compounds that ameliorate the MDS phenotype, identifying imiquimod, an agonist of TLR7 and TLR8. Imiquimod alleviates anaemia by promoting haematopoietic stem and progenitor cell expansion and erythroid differentiation, the mechanism of which is dependent on TLR7 ligation. TLR7 activation in this setting paradoxically promoted an anti-inflammatory gene signature suggesting crosstalk between pro-inflammatory pathways endogenous to Rps14 loss and TLR7 pathway activation. Finally, we show that in highly purified human bone marrow samples from anaemic patients, imiquimod leads to an increase in erythroid output from myelo-erythroid progenitors and common myeloid progenitors. Our findings have both specific implications for the development of targeted therapeutics for del (5q) MDS and wider significance identifying a potential role for TLR7 ligation in modifying anaemia.

## Introduction

Myelodysplastic syndromes (MDS) are a heterogeneous group of myeloid malignancies associated with cytopenias and evolution to acute myeloid leukaemia (AML). MDS with loss of all or part of the long arm of chromosome 5 (del(5q) MDS) is the most common subtype of MDS^1^ and, importantly, 5q loss has been shown to be an initiating event in MDS development in the majority of such cases ^2 3^.

Patients with del(5q) MDS show striking sensitivity to the immunomodulatory drug lenalidomide^4^. However, while the prognosis of del(5q) MDS is generally favourable compared to other MDS subtypes, all patients eventually fail therapy with lenalidomide with a median survival of 6 years ^5^. The only curative therapy remains stem cell transplantation, for which many older MDS patients are unfit. Thus, there remains an unmet need for novel therapies for del (5q) MDS.

Ribosomal protein of the small subunit, *RPS14* (also now known as uS11^6^), located in the critically deleted region (CDR) of chromosome 5q, is one of several genes identified as a haploinsufficient tumor suppressor gene (TSG) in del(5q) MDS ^7-12^. We and others have previously shown haploinsufficient levels of Rps14 recapitulate features of del(5q) MDS in zebrafish, mice and primary human cells ^7 13 14^.

Proposed mechanisms for Rps14-mediated haematopoietic defects include p53-dependent apoptosis and increased pro-inflammatory innate-immune signalling via toll-like receptor 4 (TLR4) resulting from an increase in translation of cognate ligands S100A8 and S100A9 ^13^. Additionally, activation of TLR-MyD88-NFKappaB(NF*κ*B) signalling in del(5q) MDS results from haploinsufficiency of other genes and microRNAs contained within the 5q CDR. Specifically, loss of miR146a leads to bone marrow failure resulting from increased expression of its target TNF receptor associated factor 6 (TRAF6), which is a key component of the TLR-MyD88-NF*κ*B signalling complex^9^. Similarly, loss of TIFAB, another candidate TSG located on 5q, also shows features of bone marrow failure in mice, with lineage^-^, Sca^+^, Kit^+^ (LSK) cells demonstrating increased inflammatory signaling and increased sensitivity to lipopolysaccharide (LPS)-mediated TLR4 signaling^11^. However, signaling through TLR4 is also required for the generation of haematopoietic stem and progenitor cells (HSPC) in both steady state and stressed haematopoiesis^15-17^. This highlights that homeostatic regulation of inflammatory signaling through TLR pathways is critical to maintain normal haematopoiesis.

The zebrafish is a highly tractable model system in which to interrogate haematopoiesis in vivo, and is amenable to high throughput in vivo small molecule screens. We therefore created a novel zebrafish model of del (5q) MDS using transcription activator-like effector nucleases (TALENs) to mutate *rps14*. We used Rps14-deficient zebrafish to identify small molecules that alleviate anaemia in this model and identified the TLR7/8 agonist imiquimod. We show that this effect of imiquimod is dependent on TLR7, and results in improved haemoglobinisation and an increase in HSPC. Importantly, we show that erythroid output is increased in both fish and primary human cells from anaemic patients in the presence of imiquimod, an effect not restricted to but enhanced by the haploinsufficiency of Rps14. Analysis of Rps14-deficient HSPC shows upregulation of negative regulators of canonical WNT signalling, which is reversed following imiquimod exposure. Furthermore we show paradoxical down-regulation of inflammatory signalling in Rps14 heterozygotes exposed to imiquimod. These data suggest crosstalk between the endogenous pro-inflammatory effects Rps14 haploinsufficiency and TLR7 activation leads to attenuation of anaemia in our models.

## Results

### Rps14 loss results in dose-dependent haematopoietic abnormalities

We have previously shown that knockdown of Rps14 using morpholinos results in anaemia, modelling the erythroid defect of del (5q) MDS^14^. To further study the effects of Rps14 loss we used TALENs to generate a stable Rps14 mutant zebrafish. This mutant carries a complex 11bp insertion-deletion creating a frameshift mutation at glutamic Acid 8 (*rps14*^*E8fs*^) (Supplemental Figure S1). To confirm this mutation specifically effects Rps14 protein levels, we used whole mount immunofluorescence in an Rps14^E8fs/-^ incross at 2dpf. This demonstrated loss of Rps14 protein in an allelic dose-dependent manner indicating that heterozygous Rps14^E8fs/+^ animals have haploinsufficient protein levels modelling that seen in del (5q) MDS (Supplemental Figure S2). Rps14^E8fs/E8fs^, herein referred to as Rps14^-/-^, show profound developmental anomalies, including reduced head size and body length, and are lethal by 5 days post fertilisation (dpf). In addition, *o*-dianisidine staining at 4dpf shows that Rps14^-/-^ embryos have markedly reduced haemoglobinisation (Figure 1C) compared to Rps14^+/+^ (Figure 1A). This phenocopies the effect observed with Rps14 morpholino knockdown, which can be rescued with Rps14 mRNA^18^. However, developmental morphology and haemoglobinisation in Rps14^+/-^ embryos was indistinguishable from their Rps14^+/+^ siblings (Figure 1A, B). In order to quantify erythroid cell numbers, we used Rps14 mutants carrying the *Tg(gata1:dsRed)* transgene which have dsRed-expressing erythroid cells. Importantly, flow cytometry of these embryos demonstrated a dose-dependent decrease in numbers of dsRed-expressing (i.e. erythroid) cells at 3dpf (Figure 1D), with a highly significant trend across genotypes (p=0.0005). This indicates that Rps14^+/-^ embryos have an anaemia associated with reduced red cell number yet are able to maintain adequate haemoglobinisation.

**Figure 1.**
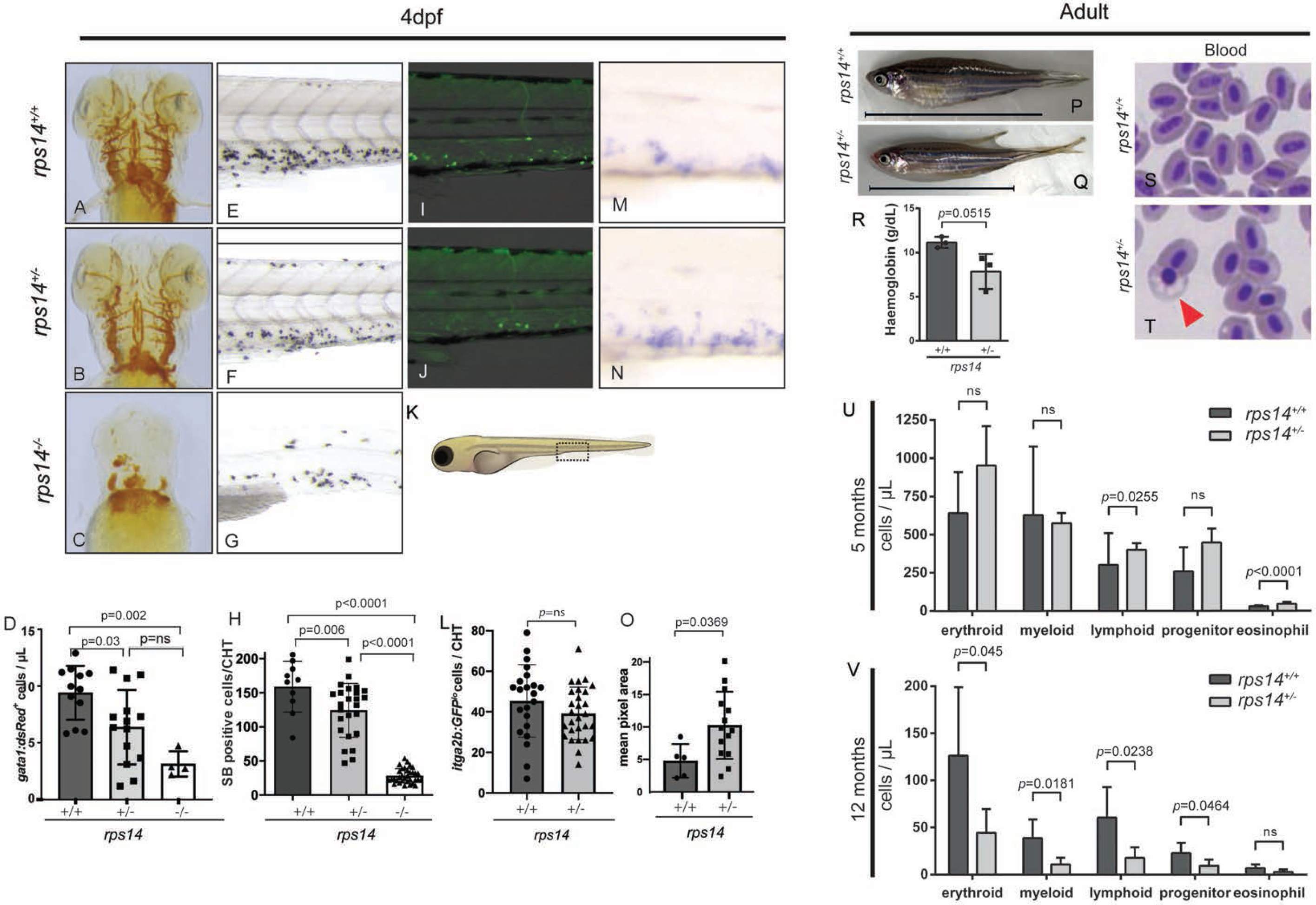
Rps14 stable mutants show dose-dependent effects on haematopoiesis in embryos. (A-C) Assessment of haemoglobinised cells using *o*-dianisidine staining. Ventral views of the head in 4dpf embryos (A-C). *Rps14*^-/-^ zebrafish show profound loss of haemoglobinised cells (C) and developmental anomalies. *Rps14*^+/-^ are indistinguishable from their WT siblings by microscopy (A, B). (D) Erythroid cell number by flow cytometry of individual *Tg(gata1:dsRed);Rps14*^*+/-*^ embryos at 3dpf. There is an allelic dose-dependent effect on dsRed-expressing cell number. (E-G) 4dpf SB-stained embryos to stain granulocytes. SB positive cell number quantified in (H). Region of CHT depicted in cartoon (K). SB staining shows an allelic dose-dependent effect of *rps14*. (I,J) Lateral views of the CHT of 4dpf *Rps14* mutant fish carrying *itga2b:GFP* transgene, labelling HSPC that reside in the CHT. (L) shows quantitation of stationary GFP^lo^ cells in the CHT. (M,N) Expression of *c-myb* by in situ hybridisation in 4dpf embryos. In contrast to *itga2b*-GFP, *c-myb* expression is increased in the CHT, quantified by median expression intensity in CHT (O)^68^. (P,Q) Lateral views of 12-month-old adult fish show a decreased size of heterozygotes. Anterior is shown to the left and dorsal upwards. Line shows the body length excluding the tail (R) shows quantification of haemoglobin concentration. (S,T) show blood smears from Rps14^+/-^ and WT siblings, red arrowhead highlighting poorly haemoglobinized erythroid cell in *rps14*^+/-^.(U) and (V) absolute cell number per microliter of different cell types in the kidney marrow of 5 and 12 months old fish respectively, showing progressive differences between heterozygous *rps14* mutants and their wild type siblings. Statistical comparison by one-way ANOVA with Tukey’s multiple comparisons test (D,H) or unpaired t-tests.

Having shown that Rps14 loss reduced erythroid cells, we next determined the impact of Rps14 loss on definitive myelopoiesis. Using sudan black (SB) to stain granulocytes at 4dpf, we identified a reduction in SB-positive granulocytes located within the caudal haematopoietic tissue (CHT) (Figure 1K) in a dose-dependent manner with allelic loss of WT Rps14 (Figure 1 E-H).

Next, we assessed the effects of Rps14 loss on HSPC. To assess this we used the transgenic zebrafish line *Tg(itga2b:GFP*) in which GFP^lo^ cells in the CHT label HSPC ^19^. HSPC quantification in the CHT of Rps14^+/-^ larvae compared to Rps14^+/+^ larva did not demonstrate any significant difference (Figure 1I-L). Rps14^-/-^ larva have virtually absent HSPC and extensive autofluorescence most likely due to dying tissue (notably all Rps14^-/-^ die by 5dpf) and were therefore not assessable. We further assessed HSPC numbers using whole mount in situ hybridisation (WISH) of *c-myb* expression in the CHT at 4dpf. By contrast to *Tg(itga2b:GFP*^*lo*^) cells, *c-myb* expression in the CHT is increased in Rps14^+/-^ compared to Rps14^+/+^ siblings (Figure 1M-O). *Itga2b*-GFP^lo^-expressing cells are some of the first to arise during definitive haematopoiesis and are restricted to HSPC and megakaryocytic lineage committed cells, while *c-myb* is expressed in a broader subset of HSPC as well as more committed myeloid lineage cells (GMP - myelocytes) (http://servers.binf.ku.dk/bloodspot/). Therefore this increase in expression in *c-myb* with normal *itga2b* and a reduction in SB stained mature myeloid cells suggests an increase in myeloid lineage restricted progenitor cells in Rps14 heterozygotes ^20^. Accordingly, these results also suggest that the reduction in lineage output observed in 4dpf embryos may result from a block in differentiation of earlier progenitor cells.

### Rps14^+/-^ adult mutants have anaemia and features of MDS

In adult fish, Rps14^+/-^ zebrafish are significantly smaller than Rps14^+/+^ (Figure 1P,Q), and have shortened body length and weight (Supplemental Figure S3A,B). Spectrophotometric analysis of haemoglobin levels showed Rps14^+/-^ adult blood contained a lower concentration of haemoglobin than that found in Rps14^+/+^ adults (Figure 1R). Furthermore, examination of haematopoietic cell morphology in May-Grünwald Giemsa (MGG) staining from Rps14^+/-^ adult fish shows poorly haemoglobinised cells in peripheral blood (Figure 1S,T) and dyserythropoiesis with nucleocytoplasmic asynchrony in the kidney marrow (Supplemental Figure S3C,D). We next used flow cytometric examination of kidney marrow (the site of adult haematopoiesis) to assess effects of Rps14 loss on haematopoiesis. At 5 months of age only an increase in the eosinophil population in *rps14*^+/-^ compared to *rps14*^+/+^ was observed by forward and side scatter examination of cell populations (Figure 1U), however we observed an increase in *Tg(itga2b:GFP*^*lo*^) cells within the progenitor cell fraction (Supplemental Figure S3E) a feature observed in murine Rps14 conditional knockouts ^21 22^. By 12 months of age the kidney marrow of Rps14^+/-^ mutants shows defects in all lineages indicating features of bone marrow failure (Figure 1V).

### Stress markedly exacerbates the haematopoietic defects in Rps14 heterozygous animals

To refine the effects of Rps14 haploinsufficiency on haematopoiesis in our model, we next exposed Rps14^+/-^ embryos to haematopoietic “stress” ^13^. First, we used cold stress, where we incubated embryos at 22°C from 10hpf for 4 days (Figure 2A). This is known to induce anaemia in zebrafish ^23^. Rps14^+/-^ embryos were significantly more anaemic following cold stress than their Rps14^+/+^ siblings (Figure 2B-D). To determine the effects of stress specifically on definitive haematopoietic cells, we investigated the effects of chemical haemolytic stress. For this, we incubated embryos in phenylhydrazine (PHZ) for 24 hours (24hpf – 48hpf) and assessed recovery of haematopoiesis (Figure 2E) ^24^. Embryos exposed to PHZ resulted in loss of all haemoglobinised erythroid cells due to apoptosis (Supplemental Figure S4). At 6dpf, WT animals had completely recovered, with *o*-dianisidine staining showing normal haemoglobinisation, while Rps14^+/-^ siblings remained markedly anaemic (Figure 2F,G quantified in 2H). We further assessed the effect of haemolytic stress on HSPC using *Tg(itga2b:GFP;rps14*^*+/-*^) transgenics. As previously, Rps14^+/-^ embryos showed no difference in *Tg(itga2b:GFP*)-HSPC numbers in the absence of PHZ. PHZ stress resulted in an increase in HSPC compared to non-stressed animals in WT, most likely in response to the induced anaemia, however Rps14^+/-^ larvae were unable to elicit this response leading to a statistically significant decrease in HSPC compared to stressed WT siblings (Figure 2I).

**Figure 2.**
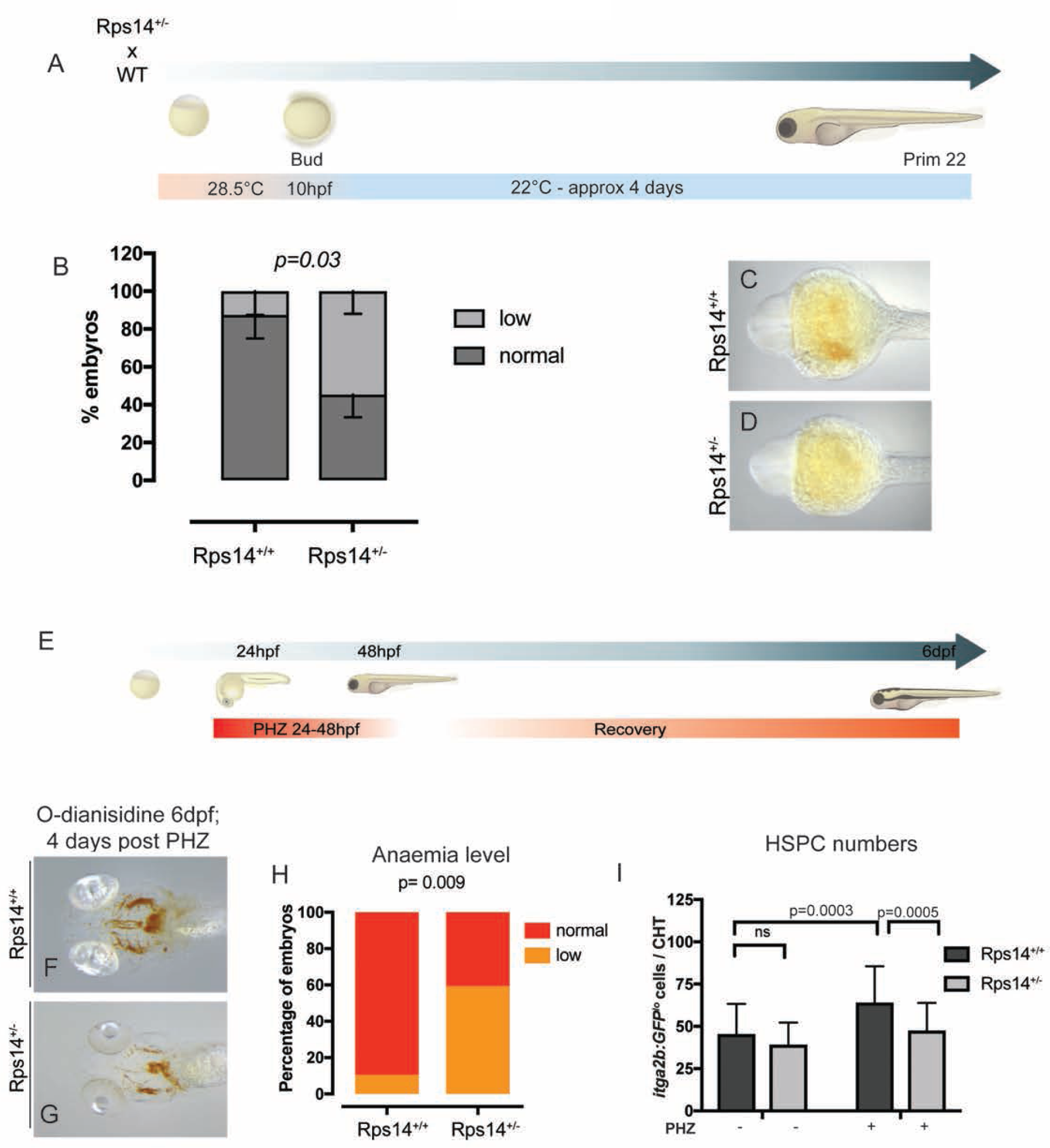
Haemolytic stress augments the haematopoietic phenotype in Rps14^+/-^ mutants, and this is rescued by imiquimod. (A) Schematic of cold stress experiment. (B) Analysis of ratio of normal:anaemic embryos in *rps14*^*+/+*^ vs *rps4*^*+/-*^ after cold stress.(C,D) Representative view (ventral, head) of Prim22 stage embryos stained with *o*-dianisidine demonstrating anaemia in *Rps14*^*+/-*^ (E) Schematic of haemolytic stress experiment (F,G) Representative views (ventral) of *rps14*^+/+^ and *rps14*^*+/-*^ siblings treated as in (E) and stained with *o*-diansidine at 6dpf. *Rps14*^*+/-*^ demonstrate a clear anaemic phenotype, quantified in (H). (I) effect of haemolytic stress on HSPC. Statistical comparisons were carried using Fisher’s exact test (B,H) or ANOVA (I).

### Small molecule screens identify imiquimod as a modifier of anaemia in Rps14-deficient embryos

To utilise our system to assess for potential novel therapeutic agents in del(5q) MDS, we developed a small molecule screen for modifiers of anaemia in Rps14-deficient embryos (Figure 3A). We used morpholino knockdown of Rps14 as the morphant phenotype is comparable to mutants, allowing us to increase the number of molecules tested ^14^. We screened the Spectrum collection (http://www.msdiscovery.com/spectrum.html) for compounds that could alleviate the characteristic morphological defects and/or anaemia resulting from Rps14 morpholino (MO) injection. Treatment of Rps14 morphants with the TLR7/8 pathway agonist imiquimod strikingly rescued the anaemia phenotype of Rps14 morphants (Figure 3D) compared to DMSO-treated negative controls (Figure 3C). 100mM L-leucine was used as a positive control (Figure 3E) and uninjected embryos as a negative control (Figure 3B).

**Figure 3.**
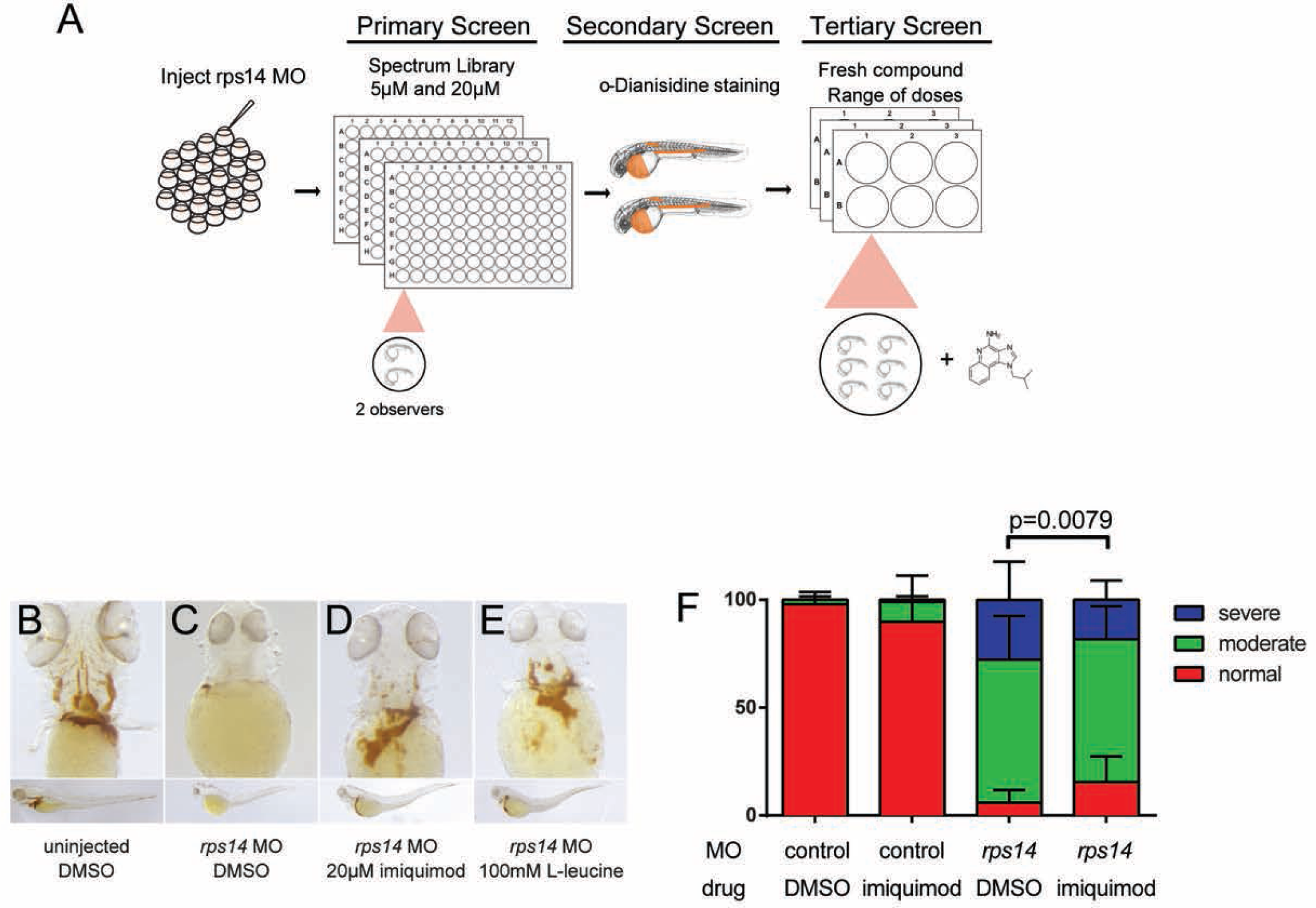
Small molecule screen for modifiers of anaemia in Rps14 deficiency identifies imiquimod. (A) Schematic of the screen design. (B-E) 4dpf *rps14* morphants and controls treated with DMSO, L-leucine or imiquimod stained for haemoglobin with *o*-dianisidine. Upper panels shows ventral views of the head, lower panels show lateral views, with anterior to the left and dorsal upwards. (F) semiquantitative analysis of the effects of imiquimod on haemoglobinization severity. Imiquimod improves the level of haemoglobinization compared to DMSO treated controls. Statistical comparisons were carried out by Fisher’s exact test (F).

We next assessed the improvement in haemoglobinisation with DMSO or 5µM imiquimod treatment by classifying each larva by the severity of their anaemia. Proportions of normal, moderate or severe anaemia in experimental groups of embryos were analysed. Figure 3F shows that imiquimod markedly decreases the proportion of larvae with severe anaemia.

### Imiquimod rescues stress-induced anaemia in Rps14 heterozygotes

To validate findings from the morphant screen, we assessed the effects of imiquimod on Rps14^+/-^ mutants exposed PHZ. As observed in Rps14 morphants, imiquimod rescued the anaemia observed in Rps14^+/-^-stressed mutants (Figure 4A-E). We also assessed the optimal dose of imiquimod and observed a dose-dependent effect of imiquimod on the proportion of anaemic animals at 6 days. The optimal dose of imiquimod with no evidence of toxicity was at 20µM imiquimod (Figure 4F).

**Figure 4.**
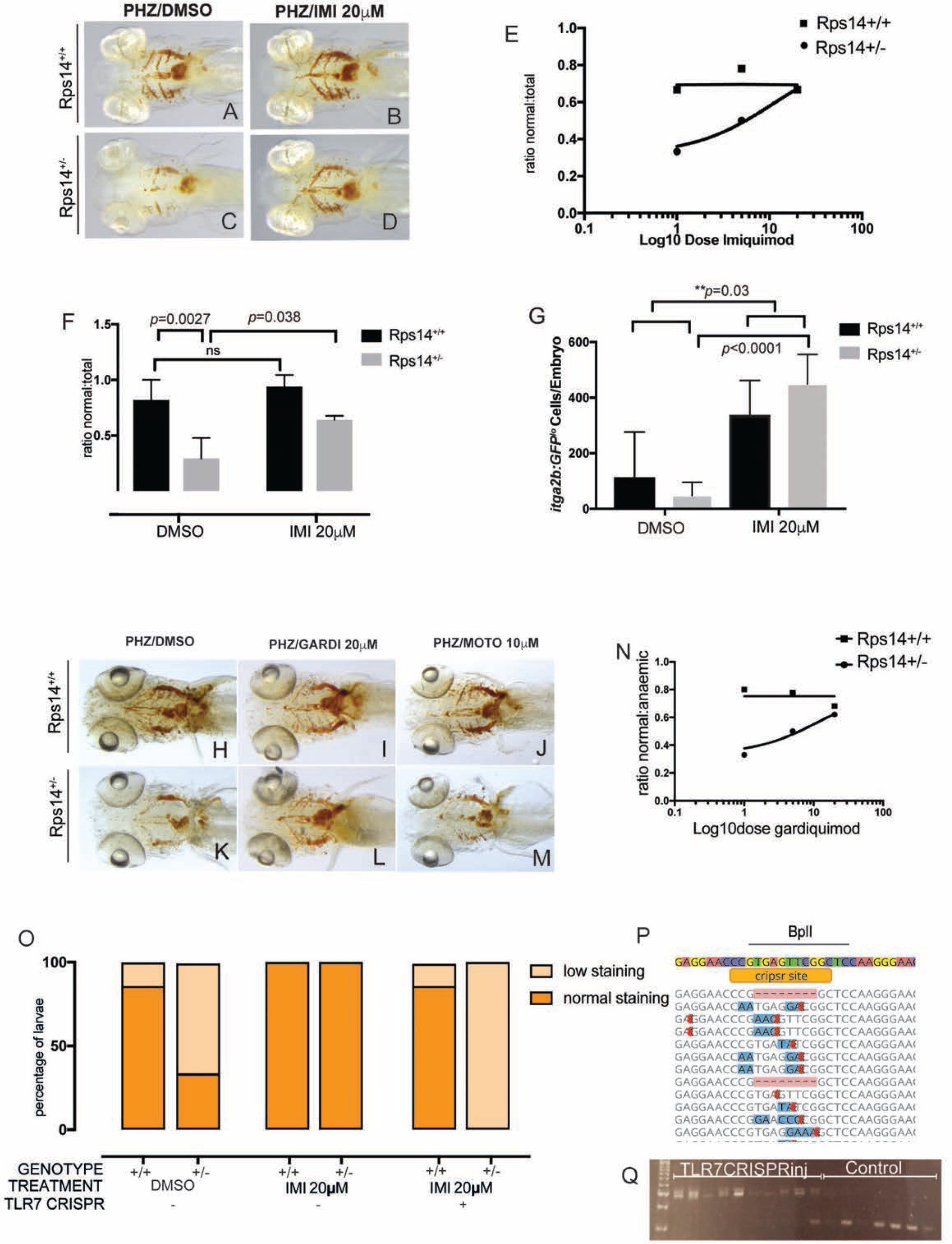
Imiquimod exerts its effect on Rps14-deficient anaemic embryos via on target activation of TLR7. (A-D) Ventral views of 6dpf embryos stained with *o*-dianisidine, treated haemolytic stress and DMSO (A and C) or imiquimod (B and D). Imiquimod rescues stress induced anaemia. Quantified as ratio of normal:total embryos across 3 experiments for 20µM (F). Effect of imiquimod analysed across dose range using non-linear regression (E). (G) Flow cytometric analysis of *Rps14*^*E8fs*^;*Tg(itga2b:GFP)* single embryos exposed to haemolytic stress and then treated with DMSO or imiquimod. Imiquimod enhances *itga2b:GFP*^*lo*^ cells and this is most marked in Rps14^+/-^ where there is a significant interaction between the drug and genotype. (H-M) Ventral views of 6dpf embryos stained with *o*-dianisidine, treated haemolytic stress and DMSO (H and K) or gardiquimod (I and L) or motolimod (J and M). Gardiquimod but not motolimod rescues the stress-induced anaemia in *Rps14*^*+/-*^ embyros. Gardiquimod rescue effect analysed across dose range using non-linear regression (N). (O) Knockout of Tlr7 abrogates rescue of anaemia by imiquimod. Tlr7 knockout validated using Miseq (P) and restriction enzyme digest with BplI (Q).

Inflammatory signalling through other TLRs has been shown to affect the emergence of HPSCs. To determine at which point during haematopoiesis imiquimod exerted its effect, and to assess the effect of TLR7 signalling on HSPC, we used *Tg(itga2b:GFP);Rps14*^*+/-*^ exposed to haemolytic stress and then treated with imiquimod or DMSO control. In PHZ stressed, control mutants, GFP^lo^ HSPC were reduced in Rps14^+/-^ compared to Rps14^+/+^ siblings. Exposure to 20µM imiquimod resulted in an increase in GFP^lo^ cells in both Rps14^+/-^ and Rps14^+/+^, however the effect was more marked in Rps14^+/-^ compared to Rps14^+/+^. A 2-way ANOVA demonstrated an interaction between the effects of the Rps14 heterozygosity and the effect of imiquimod (Figure 4G).

These findings indicate that imiquimod leads to both improved haemoglobinisation and increased HSPC numbers.

### Imiquimod impacts erythropoiesis through TLR7 ligation

Imiquimod is an imidazoquinoline. The mechanism of action of imiquimod and a number of related structures is via direct binding to TLR7 and TLR8 at equivalent residues on their dimerization interface. However, each receptor is activated in a distinct manner. TLR7 activation requires the formation of a TLR7-protomer from an inactive monomer, and the effects of imiquimod are enhanced by ssRNA (its cognate ligand) binding at a second site ^25 26^. Meanwhile TLR8 exists as a dimer in its resting state and imiquimod alters its conformation to permit downstream signalling via its TIR domain ^27 28^. Therefore to determine whether the haematopoietic effects of imiquimod we observed were through ligation of TLR7, we utilised additional small molecules that more selectively activate these receptors. To assess specificity for TLR7, we utilised the more targeted TLR7 agonist, gardiquimod. Gardiquimod treatment also rescued the anaemia associated with Rps14 heterozygosity (Figure 4H-L and N). In contrast we were unable to rescue the effects with the TLR8-specific agonist motolimod (Figure 4J, M). This indicates that the effects we observed were specific to ligation of TLR7, and that imiquimod and gardiquimod exert their effect through activation of TLR7 but not TLR8. To further confirm this we utilised a highly active TLR7 CRISPR guide RNA, confirmed to have >95% efficiency at mutating TLR7 (Figure 4P). Rps14^+/-^ x Rps14^+/+^ clutches were injected with TLR7 CRIPSR guide or control and then exposed to PHZ. Individual embryos were assessed for efficacy of knockdown using restriction fragment analysis (Figure 4Q). TLR7 crispants abrogated the imiquimod-mediated rescue of anaemia in Rps14 heterozygotes (Figure 4O). These results show that the effects of imiquimod observed occur specifically through TLR7 and do not appear to solely reflect generic activation of TLRs or off-target effects.

### Imiquimod treated Rps14^+/-^ HSPC show reversal of WNT signalling and paradoxical downregulation of inflammation and an increase in erythroid differentiation

We sought to define the mechanism by which “stress” from PHZ led to more marked anaemia in Rps14^+/-^ and whether imiquimod was alleviating this to reverse the effects. We have previously observed an increase in phosphorylation of Eif2*α* at serine 51 (Eif2*α*P) in Rps19 morphants as a putative driver of impaired translation (unpublished data). This phosphorylation event is known to occur in response to amino acid starvation, haem or endoplasmic reticulum stress and results in a global reduction in translation. We hypothesised Eif2*α*P may occur in Rps14^+/-^ under haemolytic stress due to free haem. Western blot of 6dpf whole embryo protein lysates for Eif2*α*P showed that Eif2*α*P:Total ratio was increased in Rps14^+/-^, however this was not clearly influenced by the presence of PHZ or imiquimod (Figure 5A,B). 2-way ANOVA to test whether there was an interaction between Eif2*α*P and PHZ stress or imiquimod showed no effect of either, or interaction with genotype, but demonstrated a significant effect of the Rps14 genotype alone (p=0.018). Therefore we conclude that Rps14^+/-^ results in Eif2*α*P, which contributes to defective translation in our model.

**Figure 5.**
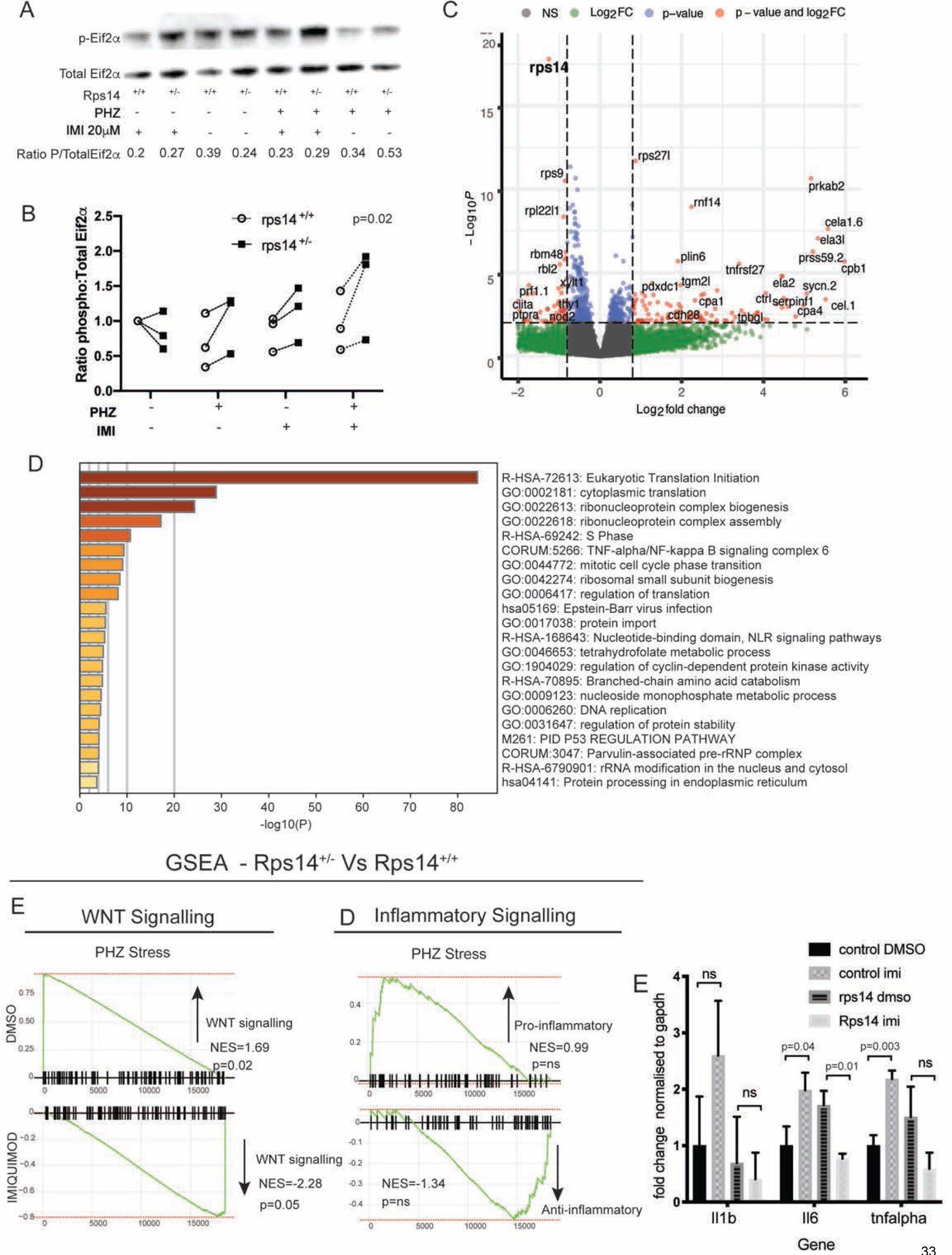
RNASeq analysis of HSPC show effects on WNT and inflammatory signalling pathways. (A) Representative western blot of p-Eif2***α*** (phosphoserine 51) and Eif2***α*** total in Rp14 mutants exposed to PHZ and/or imiquimod (B) Normalised ratio (to untreated WT) of p-Eif2***α*:**Total Eif2***α*** shown for 3 experiments (C) differential gene expression of Rps14^+/-^ vs Rps14^+/+^ shown as a volcano plot. Rps14 is highlighted in bold (D) Metascape pathway analysis of Rps14^+/-^ vs Rps14^+/+^ differentially expressed genes showing top 20 enriched pathways. (E,F) GSEA analysis comparing Rps14^+/-^ vs Rps14^+/+^ in DMSO treated vs imiquimod treated HSPC. Negative regulation of WNT signalling is enriched in Rps14^+/-^ vs Rps14^+/+^ DMSO treated HSPC and this is reversed imiquimod treated HSPC (E). Similarly inflammatory signalling is enriched in Rps14^+/-^ vs Rps14^+/+^ DMSO treated HSPC but suppressed in imiquimod-treated HSPC (F). (G) shows qPCR of TLR/NF*κ*B pathway targets using Rps14 MO or controls highlighting imiquimod rescues the effects of TLR pathway activation seen in Rps14 MO. Statistical comparisons carried out in B with 2-way ANOVA (genotype and condition)

To refine further the mechanism by which imiquimod increases HSPC and mature erythroid cells more potently in Rps14^+/-^ than in WT, we performed RNASeq analysis of HSPC with or without PHZ stress, treated with imiquimod or vehicle control. Systems level analysis of genes (visualized using Enhanced Volcano) confirm highly significant knockdown of Rps14 at the RNA level in HSPC in heterozygotes (Figure 5C)^29^. Pathway analysis of genes differentially regulated in Rps14^+/-^ compared to Rps14^+/+^ siblings using Metascape demonstrated enrichment of pathways involving ribosome biogenesis, translation, p53 pathway and TNFalpha/NF*κ*B signalling (Figure 5D), in keeping with known Rps14-associated pathways in haematopoietic development ^29 30^. The number of significantly differentially regulated genes was relatively few (35-225) between conditions, suggesting the observed phenotypes most likely arise post-transcriptionally, or through non-cell autonomous effects. To further assess changes in differentially expressed genes in order to define mediators of the effects of imiquimod, we undertook GeneSet enrichment analysis (GSEA) ^31 32^. We observed that negative regulators of canonical WNT pathway signalling were upregulated in stressed Rps14^+/-^ HSPC compared to siblings without exposure to imiquimod, but markedly downregulated following exposure to imiquimod (Figure 5E). A similar pattern was observed when we analysed inflammatory signalling signatures in stressed Rps14^+/-^ compared to siblings, with exposure to imiquimod resulting in a change from pro-to anti-inflammatory signalling (Figure 5F). To validate this finding further we conducted qPCR for downstream mediators of inflammation from TLR-signalling Il6, Tnfα and Il1b (Figure 5G). Consistent with our GSEA findings and as expected we showed that Rps14 loss results in upregulation of these inflammatory mediators, however in the presence of imiquimod these effects were reversed returning inflammatory levels to WT. These data combined suggest that Rps14^+/-^ results in pro-inflammatory signalling through the TLR-MyD88-NF*κ*B signalling complex, and treatment with imiquimod in Rps14^+/-^ paradoxically reverses this effect. This may occur via downregulation of negative regulators of canonical WNT signalling.

### Imiquimod enhances erythroid differentiation

Our data suggested that Rps14^+/-^ treated with imiquimod resulted in an increase in HSPC but also an increase in more mature erythroid cells. To assess whether this was due to a direct effect on erythroid differentiation we assessed the differential effects of Rps14^+/-^ with and without imiquimod on erythroid differentiation. GSEA showed that PHZ stress downregulates genes that promote erythroid differentiation, even at the HSPC level in *rps14*^*+/-*^ compared to heterozygotes. The addition of imiquimod reverses this effect and demonstrates a pro-erythroid differentiation signature (Supplemental Figure S5A,B). To validate this we morphologically analysed erythroid cells from sorted populations of *rps14*^*+/-*^ compared to siblings exposed to imiquimod. No clear difference in cell types was observed in this analysis (not shown), which may be due to the fact that a range of stages of differentiation were observed in both *rps14*^*+/-*^ and WT siblings. To assess effects of imiquimod on erythroid differentiation more clearly, we induced profound loss of mature erythroid cells using knockdown of Gata1 in *Tg(gata1:dsRed)* transgenic animals ^33 34^. We chose GATA1 as this master regulator of erythropoiesis has been shown as central to the mechanism of anaemia associated with RPS14 and other ribosomal proteins ^35-40^. Treatment of Gata1 morphants with imiquimod increased the numbers of circulating and static erythroid cells (supplemental Figure S5D-H and supplemental movies), and cytospin of sorted dsRed cells showed increased numbers of mature erythroid and myeloid cells (supplemental Figure S5I-K). Therefore our data support that imiquimod can not only increase HSPC numbers but promote both erythroid and myeloid differentiation on the background of anaemia associated with Rps14 or Gata1 deficiency.

### In *vitro* haematopoietic colony output is enhanced in anaemic human primary cells treated with imiquimod

Our zebrafish studies show that imiquimod increases HSPC and can alleviate anaemia by enhancing erythroid differentiation in Rps14- and Gata1-deficient anaemia. To determine if these effects were also observed in human cells we sought to analyse patient bone marrow (BM) with MDS. Unfortunately there were no patient samples with MDS del (5q) available for analysis. Therefore we utilized samples from anaemic patients with MDS or dysplastic features in their bone marrow (Table S1). HSPC output from patients with anaemia was extremely low with less than 0.02% of plated cells giving rise to any colonies (Figure 6A). This is in contrast to the non-anaemic control where 8% of plated cells gave rise to colonies (Supplemental Figure S6A) and where the overall colony output was enhanced by imiquimod.

**Figure 6.**
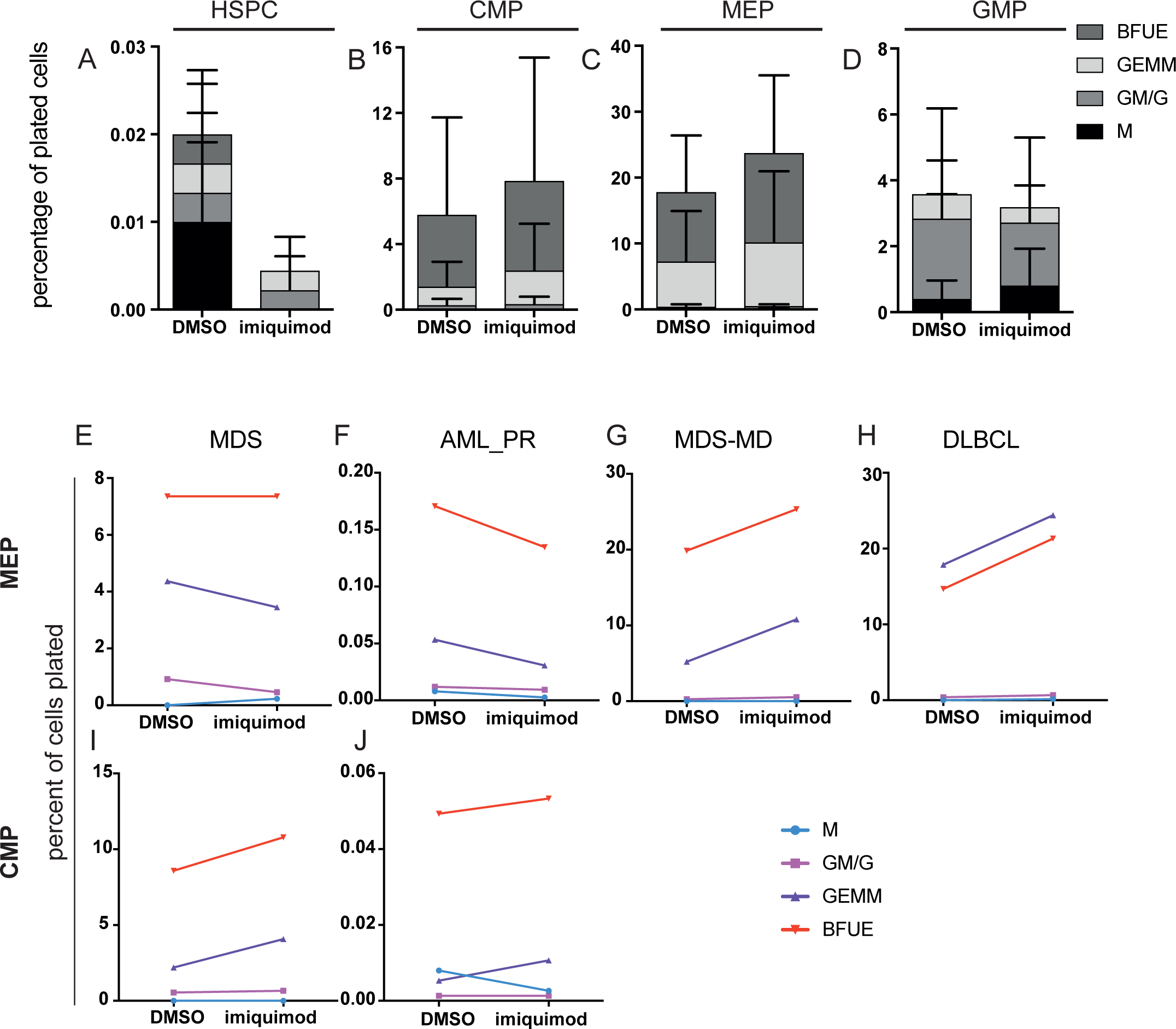
Imiquimod enhances erythroid output of human primary cells *in vitro*. (A-D) Primary cells obtained from bone marrows of anaemic patients detailed in Table S1 (n=4) were FACS-sorted into different populations. Sort purity was verified as greater than 95% for all conditions. CFU-assays were carried out in triplicate in methocult without serum. Colonies were scored at 12 days and output is shown as percentages of cells input. HSPC (A), CMPs (B), MEPs (C), and GMPs (D). (E-G) Individual patient plots highlighting the change in colony output associated with imiquimod for MEPs (E-H) and CMPs (I,J). HPSC, haematopoietic stem cells; CMP, common myeloid progenitor; MEP, megakaryocyte-erythroid progenitor; GMP, granulocyte-monocyte progenitor; BFUE, burst forming unit erythroid; GEMM, granulocyte, erythrocyte, macrophage and megakaryocyte; GM/G, granulocyte, macrophage / macrophage; M, macrophage. AML_PR, acute myeloid leukaemia in partial remission; MDS-MD, myelodysplastic syndrome with mutli-lineage dysplasia; DLBCL, diffuse large B cell lymphoma.

In both CMPs and MEPs from anaemic patients we observed an increase in colony output in the imiquimod-treated cells compared to controls (Figures 6B and 6C). For both CMPs and MEPs this reflects an increase in both colony forming units - granulocyte, erythrocyte, macrophage, megakaryocyte (CFU-GEMM) and burst forming unit-erythroid (BFU-E). In contrast there was no effect on overall colony output from GMPs, rather a lineage bias towards more mature myeloid (GM/G and M) colonies rather than GEMM (Figure 6D). To further highlight the effects at the MEP and CMP level, Figures 6E-J show the output from each patient with or without the addition of imiquimod. Taken together these data indicate that the effects of imiquimod treatment on erythropoiesis may result from effects both at the level of CMPs and MEPs in human cells. Our data also highlight that imiquimod has variable effects at different stages of lineage commitment in both the myeloid and erythroid compartment and these are not limited to Rps14 deficiency, suggesting crosstalk between mediators of inflammation may occur more widely.

## Discussion

In this study we have generated a model of del(5q) MDS using TALENS to introduce a frameshift mutation in ribosomal protein Rps14. Homozygous Rps14^-/-^ fish are embryonic lethal and developmentally phenocopy severe knockdown observed with morpholinos. Embryos deficient in Rps14 show dose-dependent defects in mature myeloid cell numbers. Erythropoiesis is similarly reduced, however, this is only macroscopically detectable in Rps14^-/-^ embryos. We have previously shown that reduction of Rps14 to approximately haploinsufficient levels using a splice-blocking MO (determined by western blot and altered splicing) results in readily detectable anaemia ^14^. This discrepancy suggests lower than haploinsufficient levels were present in the morphants. In support of this, our prior observations using MO showed that doses just lower than those utilised in our study resulted in very mild or absent effects with *o*-dianisidine staining. This supports the hypothesis that there is a threshold level of Rps14, whereby below this level the erythroid defects become more prominent.

We also show that Rps14 heterozygosity affects multiple haematopoietic lineages. GFP^lo^-expressing cells driven by the *itga2b* promoter are unchanged in steady state Rps14^+/-^ embryos compared to WT, however we observed an increase in expression of *c-myb* by WISH at 4dpf. The disparity between changes in *itga2b* and *c-myb* suggests that these markers are highlighting different subsets of cells. Indeed, *c-myb* expression, while high in HSPC, is also expressed in more committed myeloid progenitor cells, where expression of *itga2b* is markedly reduced (http://servers.binf.ku.dk/bloodspot/). We also observed that stress induced more profound alterations in haematopoiesis in Rps14^+/-^ mutants. Specifically, we observed that GFP^lo^-expressing cells in Tg(*itga2b:*GFP); Rps14^+/-^ were significantly lower in Rps14^+/-^ than WT siblings when exposed to haemolytic stress. Furthermore, Rps14^+/-^ had impaired erythroid recovery from both cold and haemolytic stressors compared to siblings. These findings suggest that Rps14 haploinsufficiency has significant effects on steady state haematopoiesis but that Rps14 also has a specific role in maintenance of stress-induced haematopoiesis. As PHZ results in stress to the haematopoietic system by haemolysing mature erythroid cells, we assessed whether Rps14-specific effects of PHZ may be mediated by stress-response pathways sensitive to free haem via Eif2*α*P. Interestingly, although we did not show an effect of PHZ on Eif2*α*P, Rps14^+/-^ mutants regardless of treatment showed an increase in Eif2*α*P compared to siblings. This suggests that a global reduction in translation arising from Eif2*α*P may contribute to the phenotypes observed.

In adults we observed age-dependent defects on haematopoiesis. Young adult zebrafish at 5 months of age show increased numbers of HPSCs marked by *itga2b*. Other committed lineages are preserved at this time point. Such HSPC defects have also been described in a conditional murine KO of Rps14 with evidence that an increase in cycling of HSPC is responsible for this increase ^13^. By 12 months of age Rps14^+/-^ animals show evidence of bone marrow failure with dysplastic features and reduction in all lineages. Thus, loss of Rps14 results in age-dependent effects on haematopoiesis. We suggest that an early developmental deficiency in HSPC is compensated for in young adults, leading to stem cell exhaustion and marrow failure in older animals.

To identify potential novel therapeutic agents for del(5q) MDS, we conducted an in *vivo* small molecule screen for compounds that could alleviate anaemia in Rps14-deficient embyros. This screen identified a striking rescue of the anaemic phenotype with the TLR7/8 agonist imiquimod. Using stress-induced anaemia as our readout, our data showed imiquimod exerts its effect on erythroid cells via TLR7 signalling. We also showed a marked effect of imiquimod on *itga2b:GFP*-expressing HPSC. Effects were observed in Rps14^+/-^ as well as Rps14^+/+^, but notably the effect in Rps14^+/-^ was more pronounced.

Pro-inflammatory signalling through TLR4 has been identified as a mechanism of anaemia in Rps14-deficient mice, and blocking this signalling pathway can rescue this effect ^13^. Rps14^+/-^ HSPC showed upregulation of TNF/NF*κ*B signalling complex in our model indicating conservation of this pathway. However TLR7 ligation by imiquimod also activates this pathway through Myd88, which results in activation of the same set of inflammatory target genes through NF*κ*B. Therefore our findings that additional activation of this pathway alleviates anaemia, appear counterintuitive. Our data support that TLR7 activation in Rps14^+/-^ HSPC paradoxically results in downregulation of inflammation. A paradigm for such crosstalk between activation of different TLRs and the effects of this on downstream signalling and homeostasis of inflammatory signalling has previously been elucidated for TLR3 and TLR7^41^. Liu *et al* demonstrated that co-activation of TLR3 and TLR7 leads to an enhanced production of some inflammatory cytokines, however, production of a number of key pathway intermediates such as TRAF6 is reduced. Co-activation of TLR7/8 and TLR9 has also been shown to synergise to reduce inflammatory output in sheep ^42^. Induction of inflammatory tolerance has also been described between TLR4 and TLR7/8 in monocytes which required microRNA 146a^43^. This suggests that while some components of the innate immune response act in concert to enhance inflammatory signalling, there are underlying mechanisms in place to halt excessive immune activation, or even specifically reduce inflammation.

The effects of imiquimod in our model were not limited to increased HSPC numbers. Using the profound differentiation arrest observed in Gata1 morphants, were demonstrated an effect of imiquimod on erythroid (as well as myeloid) differentiation. Interestingly Gata1 not only has a central role in the pathogenesis of ribosomal protein-mediated anaemias, it is also thought to be key in cytopenias associated with activation of the inflammasome in general. We did not observe a significant change in the level of Gata1 in our RNASeq datatset, however this is not surprising as this has been shown to occur post-transcriptionally ^37 38^. Notably another TLR7/8 agonist R848 has been shown to result in terminal differentiation of several myeloid leukaemia cell lines. While these authors did demonstrate effects from another more specific TLR8 agonist, they did not discount that a proportion of their phenotype occurred through activation of TLR7^44^.

When comparing Rps14^+/-^-stressed HSPC with siblings the pathway most significantly regulated in a reciprocal manner when exposed to imiquimod was WNT signalling. Specifically, stressed HSPC show a significant increase in negative regulators of the canonical WNT pathway. Regulation of HSPC specification, emergence, expansion and differentiation have all been shown to be tightly regulated, in part, by components of the WNT/β-catenin pathway ^45 46^. However, temporally, both imiquimod treatment and the effects we observe occur after specification and emergence of HSPC, suggesting in this context WNT pathway inhibition in stressed Rps14^+/-^ may impede expansion or differentiation of HSPC, and that this is alleviated by imiquimod. Furthermore crosstalk between NF*κ*B inflammatory signalling and WNT pathway activation and/or inhibition has been reported in a number of cell types (reviewed in ^47^), including evidence that prolonged TLR4-mediated inflammation suppresses WNT/β-catenin in bone resulting in apoptosis and necrosis ^48^. Our study highlights possible crosstalk between these pathways may occur but further experiments are required to validate the specific genes driving this regulation. We also assessed whether TP53 activation may be abrogated following imiquimod treatment. While TP53 target genes were upregulated as expected in the transcriptome of Rps14^+/-^ HSPC, we did not observe any change in TP53 target genes as a result of imiquimod treatment when analysing by GSEA.

TLRs, their ligands and the inflammatory consequences of signalling have been shown to differ between species^49^. For example, zebrafish have a poor response to LPS compared to mammals and carry many additional TLRs that are not present in humans^50^. However, TLR7 and TLR8 are the most highly conserved of all the TLRs and inflammatory responses to R848 are preserved in fish ^51^. Nonetheless, to establish the relevance of our findings in our zebrafish model system for humans, we utilised bone marrow cells from anaemic patients to determine the effects of imiquimod and where during lineage commitment it was affecting haematopoiesis. Our findings suggest that imiquimod can enhance erythroid output in anaemic individuals at the level of both CMP and MEP. In these experiments, very few colonies were derived from HSC of anaemic patients therefore assessment of the effects of imiquimod on HSC was not possible. One possibility is that the effects of observed in our system are non-cell autonomous via other inflammatory cells. TLR7 is expressed in zebrafish HSPC in our RNAseq data but the majority of functional effects of TLR7 signalling have been described via innate immune cells. Furthermore, we observed a larger inflammatory signalling response when analysing downstream effector genes of TLR signalling in whole embryo RNA extracts (Figure 5). Thus the cell of origin of the effects observed in our studies is not yet defined and is the focus of ongoing work.

Finally, one key unanswered question is how Rps14 deficiency and associated effects on ribosome assembly and translation leads to activation of inflammatory signalling. Murine studies define increased production of S100 proteins to be the mediators of increased inflammatory signalling via TLR4^13^. However, it is attractive to speculate, given the structural knowledge of TLR7 activation by imiquimod being enhanced by ssRNA species, that the production of aberrant rRNA species observed in Rps14 knockout cells may contribute to our observation that the effects of imiquimod are more marked in Rps14-deficient cells^7 52^. To date, no such phenomenon has been observed for endogenous RNA species outside of autoimmune disease.

In summary, we describe a model of del(5q) MDS in zebrafish amenable to *in vivo* small molecule screening and describe a novel role for the TLR 7 agonist imiquimod, identified from such a screen, in enhancing haematopoiesis and increasing the number of HSPC and their differentiation.

## Materials and methods

### Zebrafish husbandry and experimental conditions

Zebrafish (*Danio rerio*) stocks were maintained according to standard procedures in UK Home Office approved aquaria ^53^. Wild-type AB or AB/TL and transgenic strains *Tg(gata1:DsRed), Tg(itga2b:GFP)* have been previously described^19 21^. Embryos were staged according to Kimmel *et al*.^54^ and expressed in hours/days post-fertilization (hpf/dpf). All procedures complied with Home Office guidelines.

### Generation of a zebrafish *rps14* mutant line

TALENs targeting exon 2 of the zebrafish *rps14* gene were designed using Zifit targeter (http://zifit.partners.org/ZiFiT) and assembled via fast ligation-based assembly solid phase high-throughput (FLASH) as described previously^55^. TALEN mRNAs were *in vitro* transcribed using mMESSAGE mMACHINE T7 kit (Ambion, Life Technologies) and injected into one-cell stage embryos. F0 founders were identified by isolation of gametes and next-generation sequencing.

### Morpholino injection

Morpholinos (MOs) targeting the 5′UTR/ATG codon (*gata1*) or splice donor sites (*rps14*) or Gene-tools standard control were injected into 1-2 cell stage embryos at doses previously described ^14 56^.

### CRISPR/Cas9 knockdown of TLR7

A guide RNA targeting exon 2 of the zebrafish TLR7 was designed using CHOPCHOP (http://chopchop.cbu.uib.no/). This was synthesised into single synthetic modified gRNA with the sequence UUC CCU UGG AGC CGA ACU CA (TrueGuide, Thermofisher). Cas9 was in vitro transcribed from pT3TS-nCas9n (a gift from Wenbiao Chen, Addgene plasmid # 46757) using Ambion mMessage T3 kit (Ambion, Life Technologies). 300ng/µl of Cas9 was combined with the TLR7 guide at a final concentration of 5µM. This mixture was microinjected into one cell stage embyros (approximately 10pl per embryo). The level of knockdown was determined by the number of mutated reads using Miseq (Illumina) and Geneious® software or with restriction enzyme digest using BpII.

### Small molecule screen

WT zebrafish embryos injected with *rps14* MO and non-injected embryos were arrayed into 96-well plates at 24hpf (two embryos per well). Duplicate wells were treated with small molecules from the spectrum collection (Microsource Discovery Systems, Gaylordsville, USA) at 5µM and 20µM. DMSO and L-Leucine were used as negative and positive controls respectively^14^. The primary screen involved bright-field examination of morphants at 2, 3, and 4dpf. Two independent researchers scored each well for improvements in morphological development or increasing numbers of circulating erythroid cells compared to controls. Possible hits were defined as compounds scoring in a minimum of 2 wells, with at least one from each researcher. These primary candidates were re-screened in a secondary screen assessing at least 10 embryos using *o*-dianisidine staining to detect improvements in haemoglobinisation. Final hit candidate compounds were confirmed using 30 embryos per well and assessed for haemoglobinisation (5, 20, 50, 100µM) with fresh drug aliquots.

### Immunohistochemical analysis of zebrafish larvae

*O*-dianisidine staining of haemoglobin and Sudan Black (SB) staining to visualise granulocytes were carried out as previously described ^57 58^. SB positive cells were quantified from the distal end of the yolk extension to the tail tip in the caudal haematopoietic tissue (CHT; fetal liver equivalent in zebrafish).

### Induction of stress

Haemolytic stress was induced by exposure of embryos to phenylhydrazine ^59^. Embryos were incubated in 1 µg/mL PHZ from 24 hpf to 48 hpf and then washed three times. Cold stress was induced following gastrulation by placing 6 somite stage embryos at 22°C.

### Automated cell counting of *Tg(itga2b:GFP*^*lo*^) cells

Response to PHZ in *Itga2b:GFP*^*lo*^ cells was analysed using Hermes WiScan microscope (Idea Biomedical). Briefly, anaesthetised embryos were mounted into individual wells of a 96-well ZF plate (Hashimoto, Japan) and spun slowly for 20 seconds. Embryos were imaged in brightfield and GFP channel using 4 stitched images in x and 5 in z. Cells numbers were enumerated using maximal projection of GFP. GFP^lo^ cell counts in the tail were defined using WiSoft Athena software.

### Whole mount *in situ* hybridization

Whole mount *in situ* hybridizations were performed as previously described^60^.

### Probe generation

An RNA probe targeting *c-myb* was generated using one-step RT-PCR (Qiagen, Manchester, UK) using the following primers *c=myb*-F 5’-CCAAGTCAGGAAAACGCCACCTCG-3’ and *c-myb*-R 5’-GCTGTTGTTTAGCGGAGTTGGGCT-3’ and cloned into the dual promoter vector pCRII-TOPO (Life Technologies). The *pCS2:runx1* probe was a gift from Leonard Zon. Digoxigenin-labelled anti-sense RNA probes were transcribed from linearized plasmid DNA using a MEGAscript(tm) kit (Ambion, Life Technologies).

### Terminal deoxynucleotidyl transferase dUTP nick end labelling (TUNEL) staining

TUNEL staining was carried out using ApopTag ® Red *In Situ* Apoptosis Detection Kit (Millipore). Embryos were fixed in 4% paraformaldehyde and dehydrated in 100% methanol for at least 2 hours. The TUNEL staining procedure in whole embryos was undertaken as previously described ^61^.

### Human sample collection and preparation of human bone marrow for cell population sorting

Human bone marrow samples were obtained from patients undergoing routine procedures with informed consent according to UCL/UCLH Biobank for Studying Health and Disease - Haematology Project.

Fresh bone marrow was collected into EDTA and purified within 12 hours of collection by negative selection using the RosetteSep Human Hematopoietic Progenitor Cell Enrichment Cocktail according to the manufacturer’s protocol (15026, StemCell Technologies, Cambridge, UK), followed by density gradient centrifugations with Ficoll-Paque Premium (GE Healthcare Life Sciences) and SepMate(tm) tubes (StemCell Technologies).

Frozen human bone marrow mononuclear cells were thawed and washed twice with Iscove’s Modified Dulbecco’s Medium (StemCell Technologies) containing 10% fetal bovine serum (FBS). 100 μg of DNase I (07900; StemCell Technologies) per mL of cell suspension was added to prevent clumping.

Cells were washed in ice-cold PBS, 2% BSA and FcR blocking reagent (Milteyni Biotec) was added according to manufacturer’s instructions. Cells were then stained with the following antibodies for 30 minutes at 4°C: fluorescein isothiocyanate–conjugated anti-Human Hematopoietic Lineage Cocktail (contains Anti-Human CD2 RPA-2.10, Anti-Human CD3 OKT3, Anti-Human CD14 61D3, Anti-Human CD16 CB16, Anti-Human CD19 HIB19, Anti-Human CD56 CB56, Anti-Human CD235a HIR2; eBiosciences; 5 μl), phycoerythrin – conjugated anti-CD135 (4G8; BD; 10 μl), phycoerythrin-cyanine7– conjugated anti-CD45RA (HI100; BioLegend; 5 μl), Allophycocyanin-conjugated anti-CD10 (eBioCB-CALLA; eBiosciences; 5 μl), brilliant-violet-785-conjugated anti-CD38 (HIT2; BioLegend; 5 μl), and allophycocyanin-cyanine7-conjugated anti-CD34 (581; BioLegend; 2.5 μl). Hoechst 33342 (Invitrogen) was used to exclude dead cells.

Cells were sorted by flow cytometry for the following populations: HSPC: lin-, CD34+, CD38-, CD45RA-, CD10-; Granulocyte/macrophage progenitor (GMP): lin-, CD34+, CD38+, CD45RA+, CD10-, CD135+; Megakaryocyte/Erythroid progenitor (MEP): lin-, CD34+, 38+, CD45RA-, CD10-, CD135-; and Common myeloid progenitor (CMP): lin-, CD34+, CD38+, CD45RA-, CD10-, CD135+) on a FACSAria III sorter (Becton Dickenson). Flow cytometric data was analysed using FlowJo Version 10 (FlowJo LLC, Oregon).

### Colony Assays

Flow sorted cells were seeded in serum-free methylcellulose-based medium with cytokines (MethoCult SFH4436, Stemcell technologies). 5 μM imiquimod or DMSO were added to the methocult at the time of seeding in a separate location within the media prior to vortexing. Cultures were incubated at 37°C for 12 days in a humidified chamber under 7% CO_2._ CFU outputs were analysed according to the manufacturers protocol by 2 independent researchers.

### Microscopy and image processing

Microscopy was performed using a Leica M205 FA stereomicroscope using a Leica DFC310 FX camera and LAS 4.0 software. All images were processed with Fiji software version 2.0.0-rc-43/1.51f or Photoshop CS6.

### Statistical analysis

Data are presented as mean values ± SD (standard deviation). Statistical analysis was performed using GraphPad Prism version 6.0h for Mac OS X (GraphPad Software, La Jolla California USA). Specific statistical tests are shown in figure legends.

### Analysis of zebrafish by flow cytometry

Individual embryos were analysed by flow cytometry as described previously ^14^.

Adult zebrafish kidneys were dissected from terminally anaesthetised animals, dissociated in PBS with 1% FBS using a gentleMACs tissue dissociator (Milteyni) and analysed by flow cytometry.

### Cytospins

1×10^5^ cells were centrifuged onto slides, fixed with methanol and then stained with May-Grunwald-Giemsa stain. Images were taken with Hamamatsu Nanozoomer 2.0 RS

### RNASeq

Rps14^+/-^ embryos were first genotyped by tail-clipping at 3dpf and pooled 50 per condition and allowed to recover for 3 days ^62^. *Itga2b:GFP*^*lo*^ cells were sorted directly into Trizol (Invitrogen). Following RNA extraction libraries were prepared using the Smart-seq2 protocol ^63^ and sequenced on a HiSeq 2500. Data QC was conducted using FastQC and Trimmomatic and aligned to the zebrafish genome (GRCz11) using HISAT2 ^64 65^. Differential gene expression was analysed using DESeq2 and enhancedvolcano, pathway enrichment was performed using Metascape and GSEA ^30 31^ For GSEA, pathways with zebrafish gene annotations were obtained from GO2MSIG ^66^.

### Haemoglobin measurement from adult zebrafish

Adult zebrafish were venesected using heparinised 1mm glass capillary tubes as previously described ^67^. Haemoglobin concentration was determined by absorption at 540nm, using Brij® L23 solution (Sigma Aldrich; Poole, UK) and Drabkin’s reagent (Sigma Aldrich). Human haemoglobin (Sigma) was used for creation of standard curve.

### Western blotting

Protein lysates were obtained from pooling 15 6dpf genotyped embryos treated as described using. Blots were probed with Phospho-eIF2α (Ser51) (119A11) Rabbit mAb #3597 (Cell Signalling) and Total EIF2α ab26197 (Abcam). Band densities were calculated using Fiji.

### Immunofluorescence

Rabbit anti-Rps14 antibody (16683-1-AP, Proteintech) was directly conjugated to Alexa 488 using the Lightening Link® Rapid Alexa Flour 488 antibody labelling kit. Embryos were incubated overnight at 4°C in the antibody conjugate and imaged using Leica 205 with each embryo at identical zoom and fixed exposure. Genotyping was performed post-hoc and antibody intensity was quantified using Fiji pixel intensity for a defined region posterior to the midbrain-hindbrain boundary.

## Supplemental Materials

Supplemental Figure S1 relating to Figure 1 shows scheme of Rps14 and site of TALEN spacer along deleted nucleotides and predicted amino acid consequences.

Supplemental Figure S2 related to Figure 1 – Rps14^E8fs^ mutants show allele dose dependent reduction in Rps14 protein level.

Supplemental Figure S3 related to Figure 1 – Body size and weight of Rps14 mutants

Supplemental Figure S4 related to Figure 2 shows PHZ treatment results in complete loss of mature erythroid cells at 48hpf due to apoptosis of erythroid cells.

Supplemental Figure S5 related to Figure 5 shows effects of imiquimod on differentiation.

Supplemental Figure S6 related to Figure 6 shows control non-anaemic patient response to Imiquimod.

Supplemental Videos 1-4 show live *Tg(gata1:dsred)* cells in control MO and Gata1 MO injected embryos treated with Imiquimod or DMSO. Compared to the control MO, Gata1 MO leads to loss of circulating erythroid cells but accumulation of stationary very bright dsred expressing cells in the tail region.

## Authorship Contributions

EP, OP and AL designed and carried out experiments. JR, CH, AL, PD and YH carried out experiments and contributed to scientific analysis and discussion. YJ conducted the screen with assistance from MV and OP. LV and KT made TALENS targeting *rps14*. CB and SR provided essential technical assistance for human haematopoietic cell studies and analysis.

## Acknowledgements

The authors thank the CRUK-UCL flow cytometry core and UCL zebrafish facility for their assistance. EMP is supported by a CRUK Advanced Clinician Scientist Fellowship, and is a former recipient of a Wellcome-Beit Intermediate Clinical Fellowship and the Leuka John Goldman Fellowship for future science. A Medical Research Council Clinical Research Training Fellowship supports CH. OP is supported by BECAS Chile (CONICYT) and Overseas Research Scholarship (UCL). LEV contribution in this project was supported by by a FONDECYT grant (11160951) and Medical Research Council G0900994 and MR/L003775/1 to Steve W. Wilson and Gaia Gestri. CB is supported by the Swedish Research Council (no. 2015-00135) and Marie Sklodowskan Curie Actions, Cofund, Project INCA (no. 600398)

## Notes

### Competing Interest Statement

The authors have declared no competing interest.

