## Supplemental Tables and Figures for "TLR7 ligation augments haematopoiesis in Rps14 (uS11) deficiency via paradoxical suppression of inflammatory signalling and enhanced differentiation"

**Supplemental Tables, Figures and Legends**

Table S1: Patient Characteristics

| Age | Diagnosis | Disease status (%blasts; karyotype; morphology) | Hb g/l | WCC x10$^9$/l | Neuts x10$^9$/l | Plts x10$^9$/l |
|---|---|---|---|---|---|---|
| 28F | AML | 7%; Normal; dysplasia | 80 | 2.05 | 0.76 | 37 |
| 61F | Secondary MDS-EB2 | 2%; CCR; dysplasia | 86 | 1.7 | 0.62 | 42 |
| 60M | MDS-MD | 1%; CCR; dysplasia | 122 | 5.99 | 3.52 | 261 |
| 60F | DLBCL | 2%; No lymphoma; trilineage dysplasia | 95 | 8.14 | 5.94 | 322 |
| 41F | AML | 0.5%; CR 10 months post therapy end Normal; No dysplasia | 134 | 8.49 | 6 | 405 |

Abbreviations: Hb = haemoglobin; WCC = white cell count; Neuts=neutrophils; Plts = platelets; F=female; AML=acute myeloid leukaemia; MDS= myelodysplastic syndrome; MDS-EB2= myelodysplastic syndrome with excess blasts 2

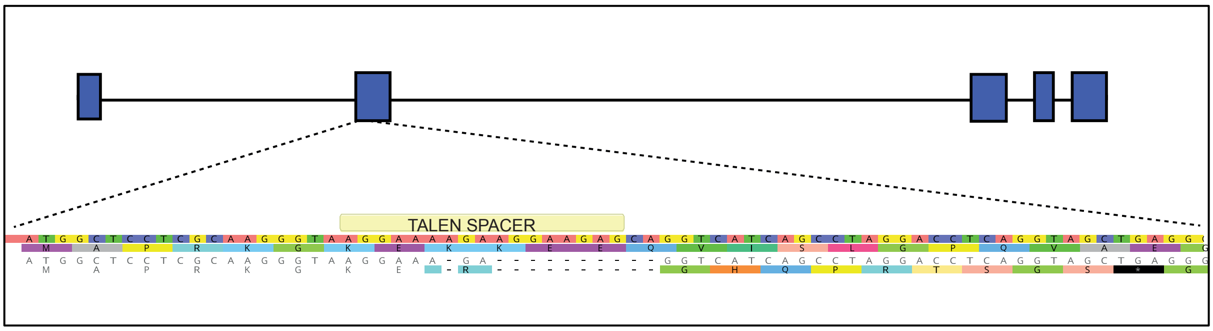


**Supplementary Figure S1. Schematic of the *rps14* gene and the localization of *rps14*$^{E8fs}$ mutation.**

WT sequence from the translational start site is shown above. Mutant sequence is shown below with predicted frameshift and truncation after 8 amino acids. Blue boxes represent *rps14* exons.

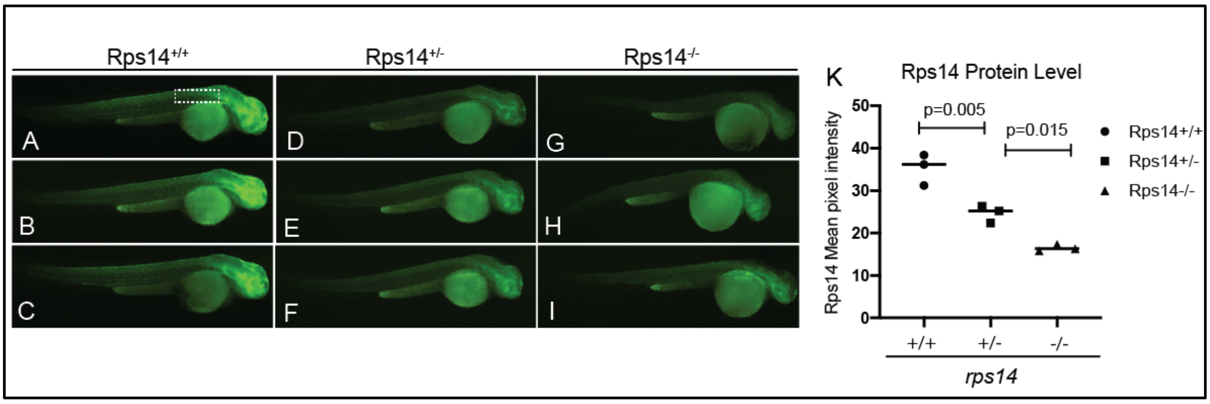


**Supplementary Figure S2. Rps14$^{e8fs}$ mutants show loss of Rps14 mutants in an allele dependent manner.**

2dpf Rps14$^{e8fs/-}$ incross clutch stained with Alexa 488 conjugated Rps14 antibody (A-C) Rps14$^{+/+}$, (D-F) Rps14$^{+/-}$, (G-H) Rps14$^{-/-}$ . Quantified using region shown in A using Fiji (K). Statistical tests carried out by one-way ANOVA with multiple comparisons.

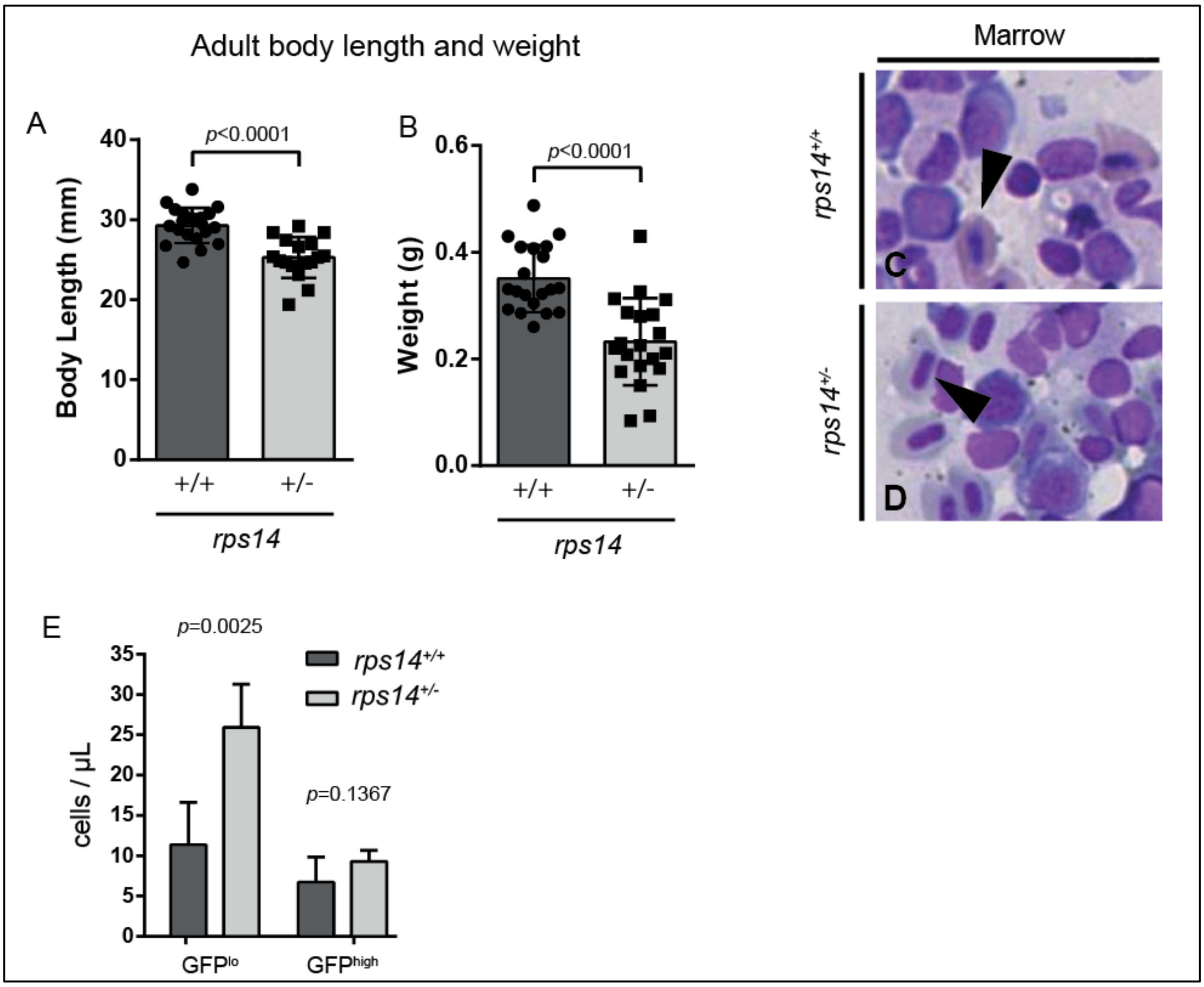


**Supplementary Figure S3. Adult Rps14$^{+/-}$ mutants are smaller than WT**

(A) Body length and (B) weight of 5-month-old Rps14$^{+/-}$ and siblings. (C,D) Representative images of kidney marrow. Black arrowheads – *rps14*$^{+/+}$ show well haemoglobinized mature erythroid cells, while *rps14*$^{+/-}$ show poorly haemoglobinized cells with nucleo-cytoplasmic asynchrony, both features of erythroid cells in MDS. Myeloid cells are morphologically normal. (E) Number of *itga2b:GFP*$^{+}$ cells in the kidney marrow of *Rps14*$^{E8fs}$*;Tg(itga2b:GFP)* 5-month-old fish. GFP$^{+}$ cells were divided according to their level of expression of GFP (GFP$^{lo}$ and GFP$^{high}$). GFP$^{lo}$ population represents progenitor cells; while GFP$^{high}$ population represents thrombocytes.

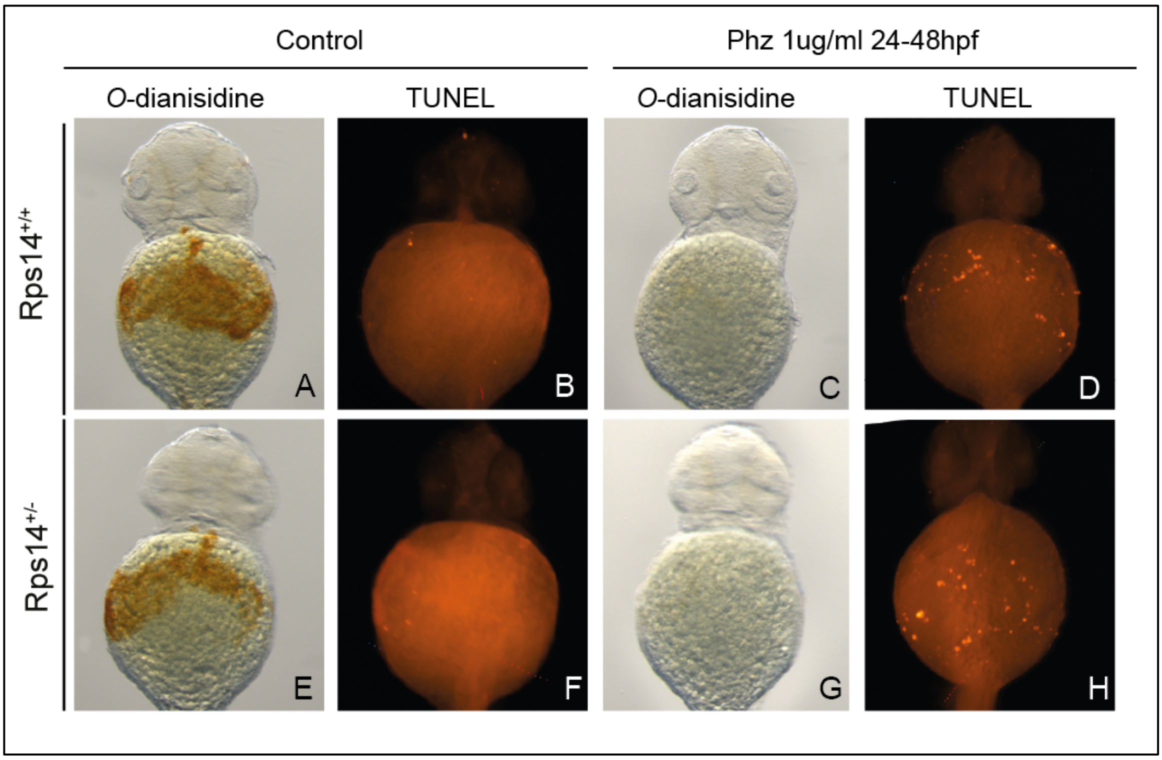


**Supplemental Figure S4 – PHZ results in cell death of all detectable mature red cells within 24 hours.**

A,C, E and G show O-dianisidine staining of 48hpf embryos treated with E3 egg water alone (A and E) or phenylhydrazine (Phz) 1µg/ml (C and G) from 24-48 hours. B, F D and H show TUNEL staining of embryos from the same treatment groups, with TUNEL positive cells highlighted in fluorescent red.

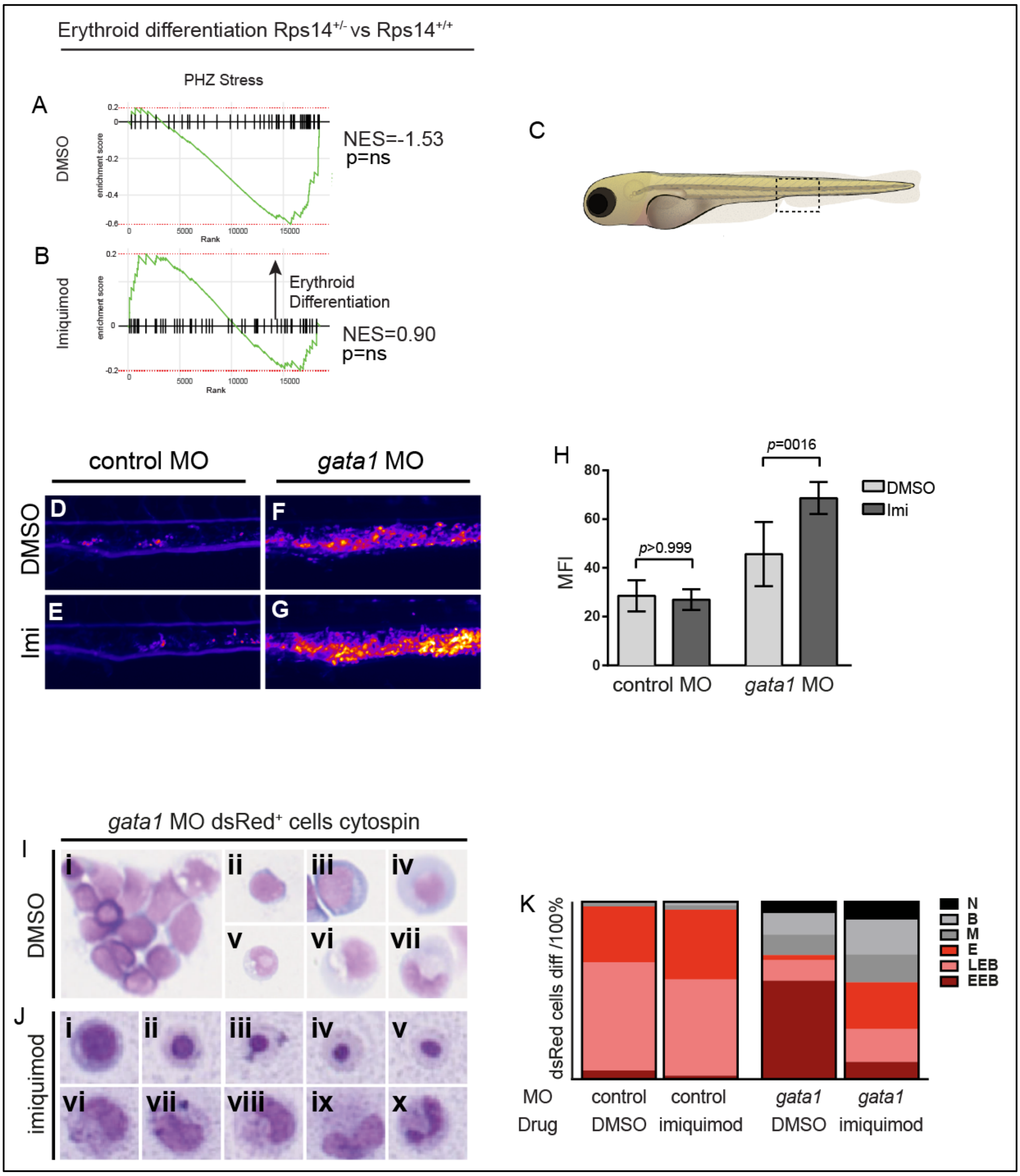


**Supplemental Figure 5. Imiquimod enhances erythroid differentiation in zebrafish embryos.**

(A,B) GSEA of Rps14$^{+/-}$ vs Rps14$^{+/+}$ for DMSO (A) and imiquimod (B) treated HSPC showing imiquimod appears to promote a differentiation gene signature where DMSO results in block in erythroid differentiation (C) Schematic of a 4dpf larva. Dotted rectangle shows the region depicted in (D-G). Representative heatmaps of fluorescence intensity of *gata1:dsRed* transgenic embryos, injected with control or *gata1* MO and treated with DMSO or 5μM imiquimod. *Gata1* morphants show marked increase in dsRed fluorescence in the CHT compared to controls (D compared to F). Treatment with imiquimod led to a further increase in fluorescence in the CHT of *gata1* morphants not observed in the control group (G compared to E). (H) Quantification of fluorescence intensity in the CHT of the morphants. (I-K) FACS sorted *gata1:dsRed*-expressing cells cytospun and stained with May-Grünwald Giemsa. Representative images of erythroid lineage cells from *gata1* morphants treated with DMSO (Ii-Iiv) or imiquimod (Ji-Jv). Imiquimod treated *gata1* morphants show increased differentiation compared to DMSO treated controls. Similarly (Iv-Ivii) shows myeloid cells in *gata1* morphants treated with DMSO compared to (Jvi-Jx) treated with imiquimod. Imiquimod increases differentiation morphologically in myeloid cells. (K) Quantification of cell types found represented as ratio of 100%. Erythroid cells are shown in different shades of red and myeloid cells in shades of grey. EEB, early erythroblast; LEB, late erythroblast; E, mature erythroid; M, myelocyte; B, band cell; N, neutrophil. Imi: imiqumod 5μM. Statistical comparisons were carried out by unpaired t-test.

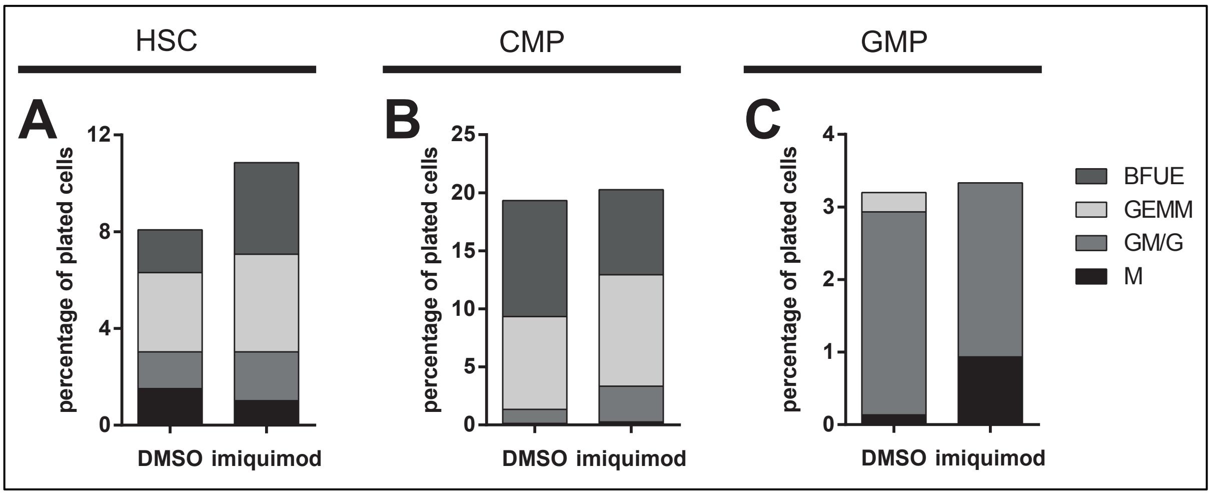


**Supplementary Figure S6.** Colony output from control AML patient in remission for 10 months with no morphological or molecular evidence of disease. Insufficient MEPs were sorted to permit plating and colony assessment.
